# MicroRNA miR-7 and miR-17-92 in POMC neurons are associated with sex-specific regulation of diet-induced obesity

**DOI:** 10.1101/366419

**Authors:** Yanxia Gao, Jiaheng Li, Zhen Zhang, Ruihan Zhang, Andrew Pollock, Tao Sun

**Affiliations:** School of Life Sciences and Biotechnology, Shanghai Jiao Tong University, Shanghai 200240, China; Center for Precision Medicine, School of Medicine and School of Biomedical Sciences, Huaqiao University, Xiamen, Fujian 361021, China; Zhiyuan College, Shanghai Jiao Tong University, Shanghai 200240, China; Department of Cell and Developmental Biology, Cornell University Weill Medical College, 1300 York Avenue, New York, NY 10065, USA

**Keywords:** miR-7, miR-17-92, obesity, POMC neurons, sex-differential gene

## Abstract

Proopiomelanocortin (POMC) neurons in the arcuate nucleus (ARC) in mammalian hypothalamus play important roles in regulating appetite, energy expenditure, and glucose and fat metabolisms. Diet-induced obesity often show sex-specific difference. But the underlying mechanisms remain unclear. Here we show that microRNA (miRNA) miR-7 and miR-17-92 are expressed in the mouse ARC, and mostly in POMC neurons. Knockdown of miR-7 and knockout of *miR-17-92* specifically in POMC neurons aggravate diet-induced obesity only in females and males, respectively. Moreover, gene expression profile analysis identifies sex-differential genes in male and female ARCs in wildtype adult mice. Interestingly, these genes that normally show low-expression in the female and male ARCs display upregulated expression in female miR-7 knockdown and male *miR-17-92* knockout mice, respectively. Our results demonstrate an important role of miRNAs in regulating sex-specific diet-induced obesity, likely through modulating expression of target genes that show sex-differential expression in the ARC of hypothalamus.

## Introduction

Hypothalamus is a conserved brain structure that regulates several basic life processes such as feeding, energy expenditure, sleep and wakefulness (Clifford & Bradford, 2015; Dietrich & Horvath, 2013; Saper et al, 2005). The arcuate nucleus (ARC) arises from cells located in the hypothalamic ventricular zone (HVZ) through progressive proliferation, lateral migration and differentiation (Mcclellan et al, 2008; Shimogori et al, 2010; Toda et al, 2017). The ARC consists of two major types of neurons, orexigenic neurons that produce neuropeptide Y (NPY) and agouti-related protein (AgRP) (Joly et al, 2010; Mercer et al, 2011), and anorexigenic neurons that produce proopiomelanocortin (POMC) and cocaine- and amphetamine-related transcript (CART) (Gao & Sun, 2016; Greenman et al, 2013). POMC neurons are identified as early as embryonic day 10.5 (E10.5) in the mouse brain and their functional circuits become mature by two-weeks-old (Coupe & Bouret, 2013; Toda et al, 2017). Studies have shown that POMC neurons function in decreasing food intake (Clifford & Bradford, 2015), regulating glucose metabolism (Berglund et al, 2012; Parton et al, 2007; Smith et al, 2015), and promoting fat burning (Dodd et al, 2015).

Interestingly, biological functions of hypothalamus show sex-specific preference (Swaab et al, 2001). For example, restricting POMC expression in the 5-hydroxytryptamine (serotonin) receptor 2C positive cells in male and female mice, only male mice show hyperinsulinemia and higher capacity of fat burning (Burke et al, 2016). Knockout of *Pten* increases body weight of male mice under the normal diet, while it increases body weight of female mice under high fat diet (Plum et al, 2007). The underlying mechanisms of sex-specific preference of hypothalamic functions are still unclear.

MicroRNAs (miRNA) are about 22 nucleotides small RNAs that normally affect mRNA stability or block mRNA translation by complementary sequence binding to the 3’ untranslated region (3’UTR) of their targets (Bartel, 2009; Lai, 2002; Pasquinelli, 2012). Studies have shown important role of miRNAs in neural development and function (Bian et al, 2013b; Gao, 2008; Rajman & Schratt, 2017). Some miRNAs have been found to show enriched expression in hypothalamus, for example miR-7, let-7 and miR-124 (Bak et al, 2008). Expression levels of some miRNAs such as miR-200a and miR-383a are increased in hypothalamus in obesity models (Crepin et al, 2014; Derghal et al, 2015). Moreover, expression of miR-103 in hypothalamus can protect mice from obtaining obesity (Vinnikov et al, 2014), while deletion of *Dicer*, an miRNA processing enzyme, in POMC neurons results in obesity (Schneeberger et al, 2012; Vinnikov et al, 2014). These studies imply that miRNAs in hypothalamus are involved in regulating functions of POMC neurons. However, it is unclear which specific miRNAs play a role in controlling development and function of POMC neurons.

Here we show that miR-7 and miR-17-92 are expressed in POMC neurons in the mouse brain. Knockdown of miR-7 in POMC neurons aggravates diet-induced obesity in females not males. And *miR-17-92* knockout in POMC neurons causes diet-induced obesity in males. Upregulation of female-low expression genes in miR-7 knockdown mice and male-low expression genes in *miR-17-92* knockout contributes to sex-specific obesity. Our results demonstrate roles of specific miRNAs such as miR-7 and miR-17-92 in regulating POMC neuron development and function, in particular in a sex-specific manner.

## Materials and Methods

### Experimental Animals

All experimental mice were housed in the animal facility of Shanghai Jiao Tong University. All animal experiments were approved by the animal ethics committee of Shanghai Jiao Tong University.

To conditionally knock out miR-17-92 in the POMC neurons, *miR-17-92^flox/flox^* mice were bred with *Pomc-Cre* mice to generate *Pomc-Cre;miR-17-92^flox/flox^*, called *miR-17-92* KO mice (Bian et al, 2013a; Jin et al, 2016). To conditionally knock down miR-7 in the POMC neurons, *miR-7* sponge transgenic mice were bred with *Pomc-Cre* mice to generate *Pomc-Cre;miR-7-sponge*, called *miR-7-sp* mice (Pollock et al, 2014).

To visualize POMC neurons, *tdTomato* reporter transgenic mice were bred with *Pomc-Cre* mice to generate *Pomc-Cre;tdTomato* mice, in which the *tdTomato* reporter gene is activated in the POMC neurons. Moreover, the *Pomc-Cre;miR-7-sponge;tdTomato* (called *miR-7-sp^tdTomato^*) mice and control *Pomc-Cre;tdTomato* (named *Ctrl^tdTomato^*) mice were generated by crossing homozygous *tdTomato* transgenic mice with *Pomc-Cre*;*miR-7*-*sponge* mice. Moreover, *Pomc-Cre;miR-17-92^flox/flox^;tdTomato* (called *miR-17-92* KO*^tdTomato^*) mice and *Pomc-Cre;miR-17-92^flox/+^;tdTomato* (named *Ctrl^tdTomato^*) mice were generated by breeding *miR-17-92^flox^/^flox^* mice with *Pomc-Cre;miR-17-92^flox/+^ ;tdTomato* mice.

### Glucose metabolism

The intraperitoneal glucose tolerance test (IPGTT) measures the clearance of injected glucose. It is used to measure glucose metabolism that is related to human diseases such as diabetes or metabolic syndrome (Latreille et al, 2014; Parton et al, 2007). Briefly, mice were fasted for consecutive 16 hours and subsequently the blood glucose level was measured as the baseline. D-glucose was injected intraperitoneally with the dose of 2g/kg body weight in mice at 16 weeks old, and with the dose of 1g/kg body weight in mice at 35 weeks old. The blood glucose level was then measured 15, 30, 45, 60, 90 and 120 minutes after D-glucose injection with an ACCU-CHEK active glucometer (Roche).

### Tissues preparation

For antibody staining, mouse brains at E15.5 and P0 stages were collected directly, and postnatal brains were collected after perfusion of phosphate-buffered saline (PBS) and 4% paraformaldehyde in 0.1M PBS (PFA). All brains were post-fixed with 4% PFA overnight and dehydrated in 30% (w/v) sucrose for a few hours (E15.5) or a few days (P0, P14 or adult brain) till brains sunk to the bottom of tubes. Dehydrated tissues embedded in O.C.T (Sakura) were sectioned (10-15 μm) by a cryostat. According to tdTomato signals, coronal sections of ARC nucleus were collected under the fluorescence stereomicroscope (Leica).

For RNA extraction, brains were immediately removed after anesthetization by 3.5% chloral hydrate. Coronal brain sections including ARC were collected by a 0.5mm mouse stainless steel brain matrices (RWD, Shenzhen), and fresh ARC tissues were separated under the microscope. The whole collecting process was performed in cold 0.1M PBS buffer to suppress ribonuclease activity.

### Immunofluorescent staining

Slides were dried up in room temperature and then boiled in the boiling buffer (800 ml milli-Q water, 4 ml 1M Tris pH8, 1.6 ml 0.5 EDTA) for antigen recovery. Then brain sections were blocked in 150 μl blocking buffer (0.1M PBS; 10% NGS and 0.1% Tween 20), the first antibody rabbit anti-RFP (1:300) was incubated overnight and signals were visualized using the secondary antibody Alexa-Fluor-647 (1:300) with incubation for 1.5 hours. After nucleic DNA staining by 4’,6-diamidino-2-phenylindole (DAPI), slices were mounted with fluorescent anti-fade mounting medium (Dako). Images were captured using the TCS SP8 confocal microscope (Leica).

### Quantitative real-time reverse transcription PCR (qRT-PCR)

Tissues were homogenized by the TRIZOL reagent (Invitrogen), and total RNA was separated by chloroform and precipitated by isopropanol. After washing it with 75% ethanol, the RNA pallet was dissolved in RNase-free water. And then 500ng total RNA was reversely transcribed into cDNA by reverse transcriptase (Takara). Quantitative PCR was performed in the qPCR machine (Bio-Rad) using SYBR green Mix (Takara). The primers used were shown in Table S1. The relative mRNA level was calculated by the method 2^−ΔΔT^.

### Statistical analysis

For two groups comparison, independent Student’s t-tests were performed after equality of variances were assessed by Levene’s test, and ANOVA analysis were used for multiple comparisons in which least significant difference (LSD) tests were conducted when variances were equal, otherwise Tamhane’s T2 tests were performed. All tests were two-tailed and *P* values <0.05 were considered statistically significant. Values were presented as mean±standard error mean.

## Results

### miR-7 is expressed in POMC neurons in developing mouse brains

When examining miRNA function in cortical development, we found that some miRNAs such as miR-7 are also expressed in hypothalamus (Pollock et al, 2014). To verify miR-7 expression in the arcuate nucleus (ARC), we performed *in situ* hybridization using locked nucleic acid (LNA) probes of mature miR-7-5p. miR-7 expression was detected in E15.5 brains, including the ARC (Figure 1A). In postnatal day 0 (P0) brains, miR-7 expression was slightly increased in the ARC compared to that in E15.5 brains (Figure 1B). miR-7 expression was maintained in the P14 ARC (Figure 1C). Moreover, in serial coronal sections collected from the anterior to posterior brain regions of adult mice (4 months old), miR-7 displayed higher expression in the medial ARC than the anterior and posterior (Figure S1).

**Figure 1.**
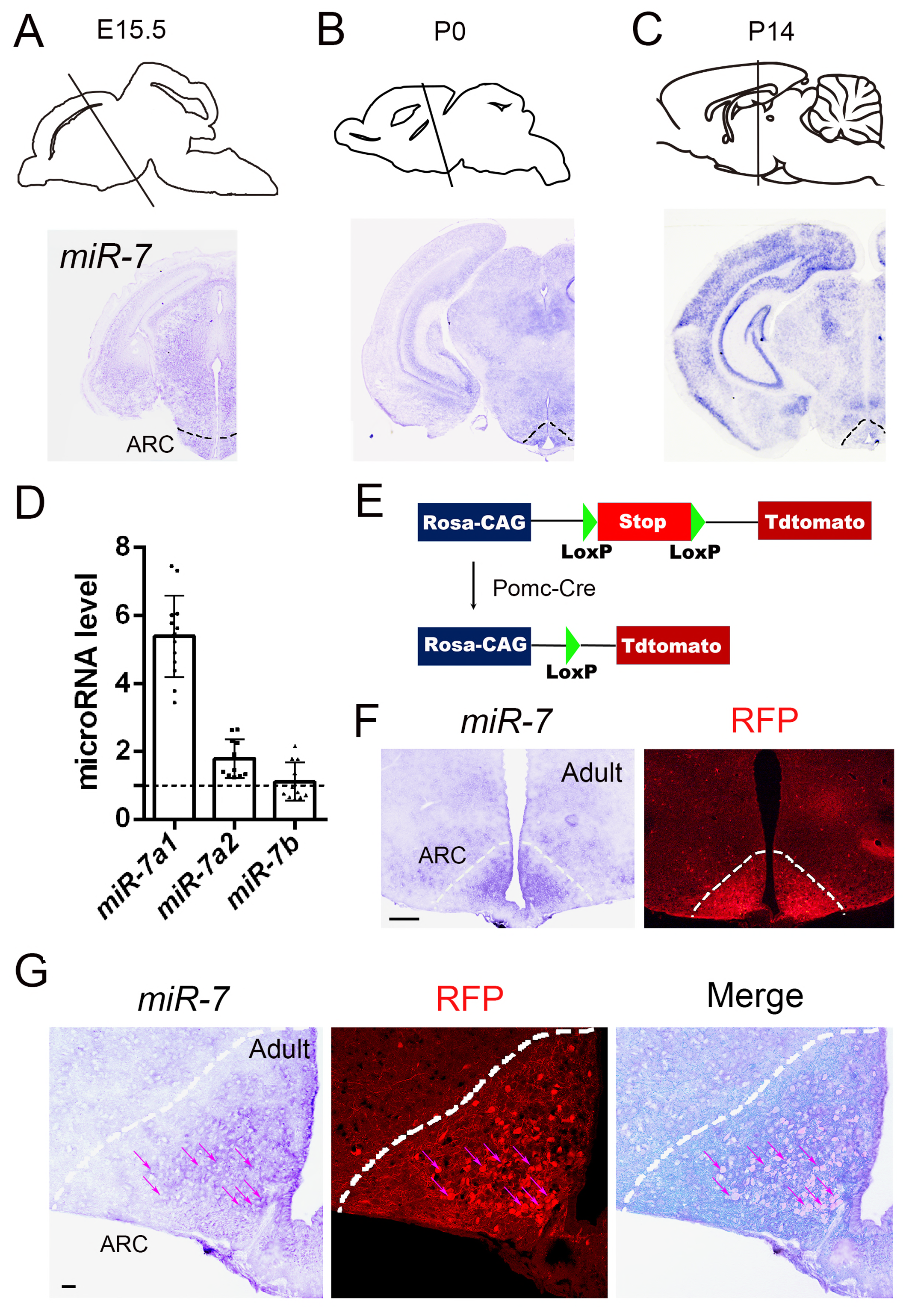
*miR-7* expression in POMC neurons in the mouse brain. A-C miR-7 expression patterns in the arcuate nucleus (ARC) in E15.5, P0 and P14 brains detected by *in situ* hybridization. Dashed lines illustrate the ARC. D Relative expression levels of members of miR-7 family in the adult ARC detected by quantitative real time reverse transcription PCR (qRT-PCR). miR-7a1 displayed the highest expression level. n=11-13 brains in each group, values shown are means ± s.e.m. E Diagram of generation of *Pomc-Cre;tdTomato* mice to visualize POMC neurons. F Images of miR-7 *in situ* hybridization (left) and RFP immunohistochemistry (middle). Dashed lines illustrate the ARC. Scale Bar: 200μm. G miR-7 was expressed in most POMC neurons in the ARC. Images of miR-7 *in situ* hybridization (left), RFP immunohistochemistry (middle) and merge of both (right). Dashed lines illustrate the ARC, and arrows mark neurons with co-expression of miR-7 and RFP. Scale Bar: 75μm.

Mature miR-7 consists of 3 family members with identical seed sequence: miR-7a1, miR-7a2 and miR-7b (Pollock et al, 2014). To distinguish expression levels of 3 miR-7 members in the adult ARC, we conducted real time quantitative RT-PCR (qRT-PCR). miR-7a1 displayed the highest expression level among all 3 members (Figure 1D). These results suggest that miR-7a1 might play a major biological role in the miR-7 family.

Considering POMC neurons mainly reside in the ARC (Padilla et al, 2010), we next examined whether miR-7 is expressed in POMC neurons. To visualize POMC neurons, *tdTomato* reporter transgenic mice were bred with *Pomc-Cre* mice to generate the *Pomc-Cre;tdTomato* line, in which most of POMC neurons will be visible under a fluorescent microscope due to specific *Pomc-Cre* activity (Madisen et al, 2010; Padilla et al, 2012) (Figure 1E). In adult brain sections of *Pomc-Cre;tdTomato* mice, *in situ* hybridization was conducted using the miR-7-5p LNA probe, followed by immunohistochemistry using anti-RFP antibodies to label POMC neurons (Figure 1F and Figure S1). Images of RFP signals and miR-7 expression in the ARC were collected under the fluorescent and bright fields, respectively, and overlaid. A large number of cells (83.8% co-marked cells among RFP positive cells, based on 3 individual brain sections) in the ARC were co-marked with miR-7 probes and RFP (Figure 1G). These results suggest that miR-7 is mostly expressed in adult POMC neurons.

### Knockdown of miR-7 results in obesity in female not male mice under high fat diet

Given the role of POMC neurons in food intake, energy expenditure and glucose metabolism (Toda et al, 2017), we examined whether miR-7 is involved in regulating these processes. To knock down miR-7 expression, the miRNA sponge technology is applied (Pollock et al, 2014). *miR-7-sponge* transgenic mice were bred with *Pomc-Cre* mice to generate *miR-7-sp* mice in which miR-7 is specifically knocked down in POMC neurons (Pollock et al, 2014). Wildtype and/or *miR-7-sponge* transgenic mice without *Pomc-Cre* were treated as control mice.

We first examined the number of POMC neurons after miR-7 knockdown. To visualize POMC neurons in *miR-7-sp* mice, homozygous *tdTomato* mice were crossed with *miR-7-sp* mice to generate *Pomc-Cre;miR-7-sponge;tdTomato* mice (called *miR-7-sp^tdTomato^*), and *Pomc-Cre;tdTomato* control mice (named *Ctrl^tdTomato^*). Immunohistochemistry using anti-RFP antibodies to label POMC neurons was conducted in sections collected from the most anterior to the most posterior ARC in P0 brains. The total numbers of RFP^+^ neurons were counted in every 5 sections from the anterior to posterior ARC. The numbers of RFP^+^ POMC neurons didn’t show significant difference between *miR-7-sp^tdTomato^* and *Ctrl^tdTomato^* mice at P0 (Figure S2A, B). These results indicate that knockdown of miR-7 does not affect POMC neuron development.

We next examined whether knockdown of miR-7 in POMC neurons affects the body weight of mice under normal chow diet (NCD), and we found no significant changes of body weights in control and *miR-7-sp* mice (Figure S3A). Interestingly, when high fat diet (HFD) was given, even though the amount of food intake was similar between males and females in control and *miR-7-sp* mice, while male mice did not show changes in body weights, *miR-7-sp* female mice displayed significantly higher weights than control females (Figure 2A-C and Figure S3B-D).

**Figure 2.**
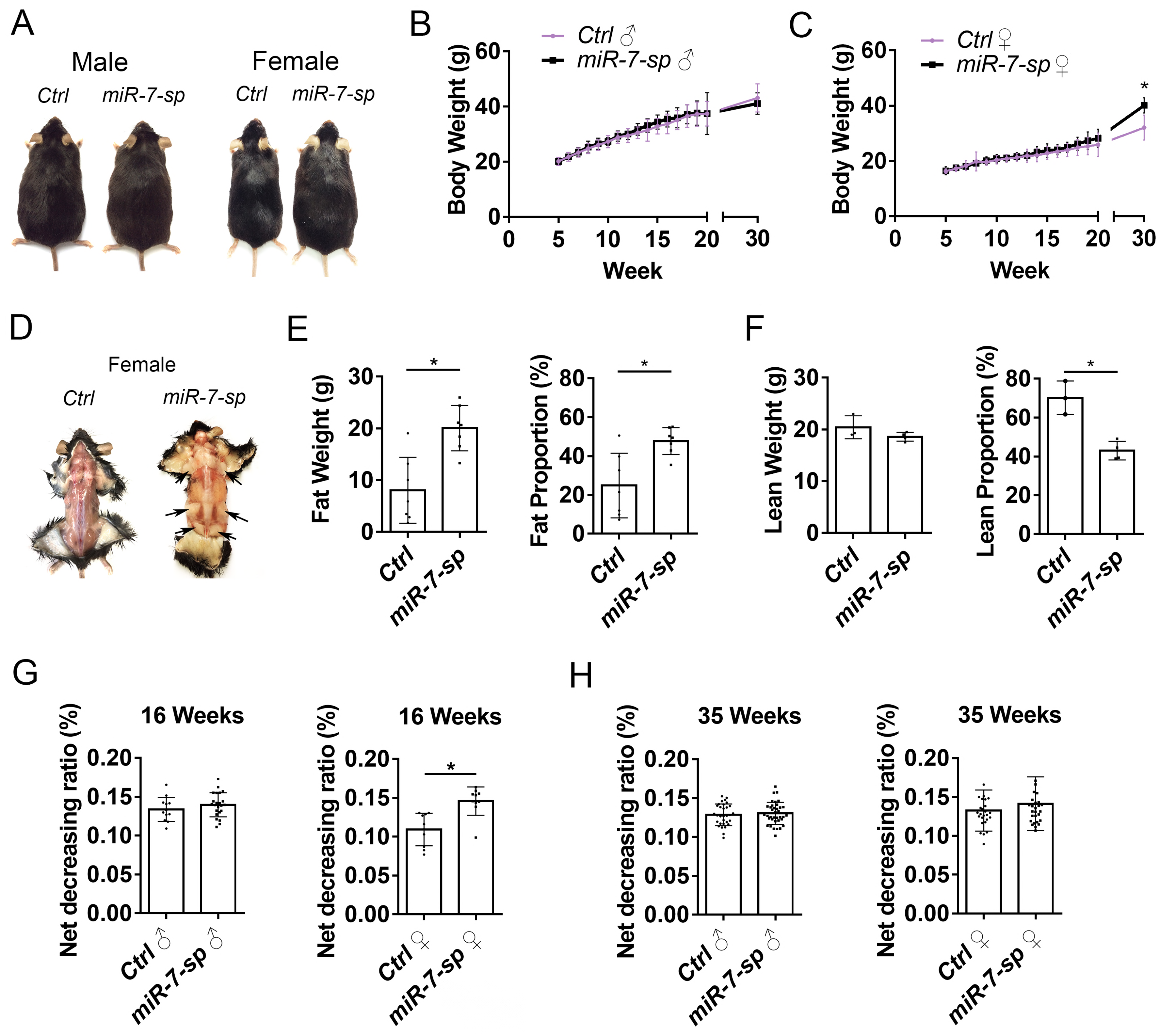
Knockdown of miR-7 results in obesity in female not male mice under high fat diet. A Images of male and female miR-7-sp and control (Ctrl) mice (35-weeks-old) under high fat diet (HFD). B, C Body weight of male (B) and female (C) miR-7-sp and control mice from 5 to 30 weeks old after treating with HFD. Compared to the same sex control mice, body weights of female miR-7-sp mice were increased significantly. n=10-28 mice at each time point. D Fat tissues of female miR-7-sp and control mice under high fat diet at 35 weeks old. E, F Body composition of HFD-fed female mice at 35 weeks old. Fat mass and fat proportion (fat weight/body weight) of female miR-7-sp mice were significantly higher than those of controls (E). Lean mass had no difference between female miR-7-sp and controls, but lean proportion of miR-7-sp mice was significantly lower than that of controls (F). n=3-7 mice in each group. G, H Changes in body weights after fasting for 48 hours at 16 weeks (G) and 35 weeks (H) under normal chow diet. The ratio of body weight decrease in female miR-7-sp mice was more than controls at 16 weeks old. n=9-22 mice in each group. Values shown are means ± s.e.m. *: P<0.05, Student’s t-test.

Moreover, body composition in female mice was measured by magnetic resonance imaging (MRI) analysis (Figure 2D). The fat weight and fat proportion were significantly increased in *miR-7-sp* females, compared to control females (Figure 2E). Consequently, the lean proportion was greatly decreased in *miR-7-sp* females, compared to control females, even though the lean weight did not change (Figure 2F). These results indicate that *miR-7* knockdown specifically aggravates obesity in females not in males under high fat diet.

Furthermore, under the normal chow diet condition, *miR-7-sp* and control mice were challenged by being fasted for 48 hours at 16 and 35 weeks old. While there was no difference in male mice, *miR-7-sp* female mice showed a higher ratio of weight decrease than control females at 16 weeks old (Figure 2G). However, the weight decrease in *miR-7-sp* female mice was recovered at 35 weeks old (Figure 2H). These data suggest that knockdown of miR-7 transiently affects response to food fasting in female not male mice.

### Knockdown of miR-7 affects glucose metabolism

The intraperitoneal glucose tolerance test (IPGTT) measures blood glucose clearance to assess glucose metabolism (Latreille et al, 2014; Parton et al, 2007). The IPGTT test was first conducted in *miR-7-sp* and control mice under normal chow diet. Interestingly, after 15, 30, 45, and 60 minutes of glucose injection, 18 weeks old female *miR-7-sp* mice showed lower glucose levels than the same aged controls, while male *miR-7-sp* mice displayed compatible levels with controls (Figure S3E). However, neither female nor male *miR-7-sp* mice showed difference at 35 weeks old (Figure S3F). These results suggest that knockdown of miR-7 results in periodic stronger abilities to glucose clearance in female not male mice.

Furthermore, the IPGTT test was conducted in *miR-7-sp* and control mice under high fat diet. Opposite to the normal chow diet conditions, after 30, 60, and 90 minutes of glucose injection, 18 weeks old male *miR-7-sp* mice showed lower glucose levels than the same aged controls, while female *miR-7-sp* mice displayed compatible levels with controls (Figure S3G). Neither female nor male *miR-7-sp* mice showed difference at 35 weeks old (Figure S3H). These data suggest that under high fat diet, knockdown of miR-7 causes periodic stronger abilities to glucose clearance in male not female mice. These results also hint that miR-7 regulates glucose metabolism in a sex-specific manner.

### MicroRNA miR-17-92 cluster is expressed in POMC neurons

Our previous study has shown expression of miR-17-92 cluster in the cortex and hypothalamus (Bian et al, 2013a; Jin et al, 2016). The miR-17-92 cluster consists of 4 subfamilies: miR-17 and miR-20a; miR-19a and miR-19b; miR-92a; and miR-18a (Bian et al, 2013a). Because miR-18 normally had low copy number of expression (Bian et al, 2013a; Jin et al, 2016), we validated expression patterns of miR-17, miR-19a and miR-92a using *in situ* hybridization. All 3 miRNAs were expressed in the ventral hypothalamus at E15.5 and P0, particularly in the ARC (Figure S4A and B). In serial coronal sections collected from the anterior to posterior brain regions of adult mice, all 3 miRNAs displayed higher expression in the posterior and medial ARC than the anterior ARC (Figure 3A and Figure S4C). We next examined expression levels of 3 miRNAs in the adult ARC using qRT-PCR. miR-92a displayed the highest expression level among all 3 members (Figure 3B).

**Figure 3.**
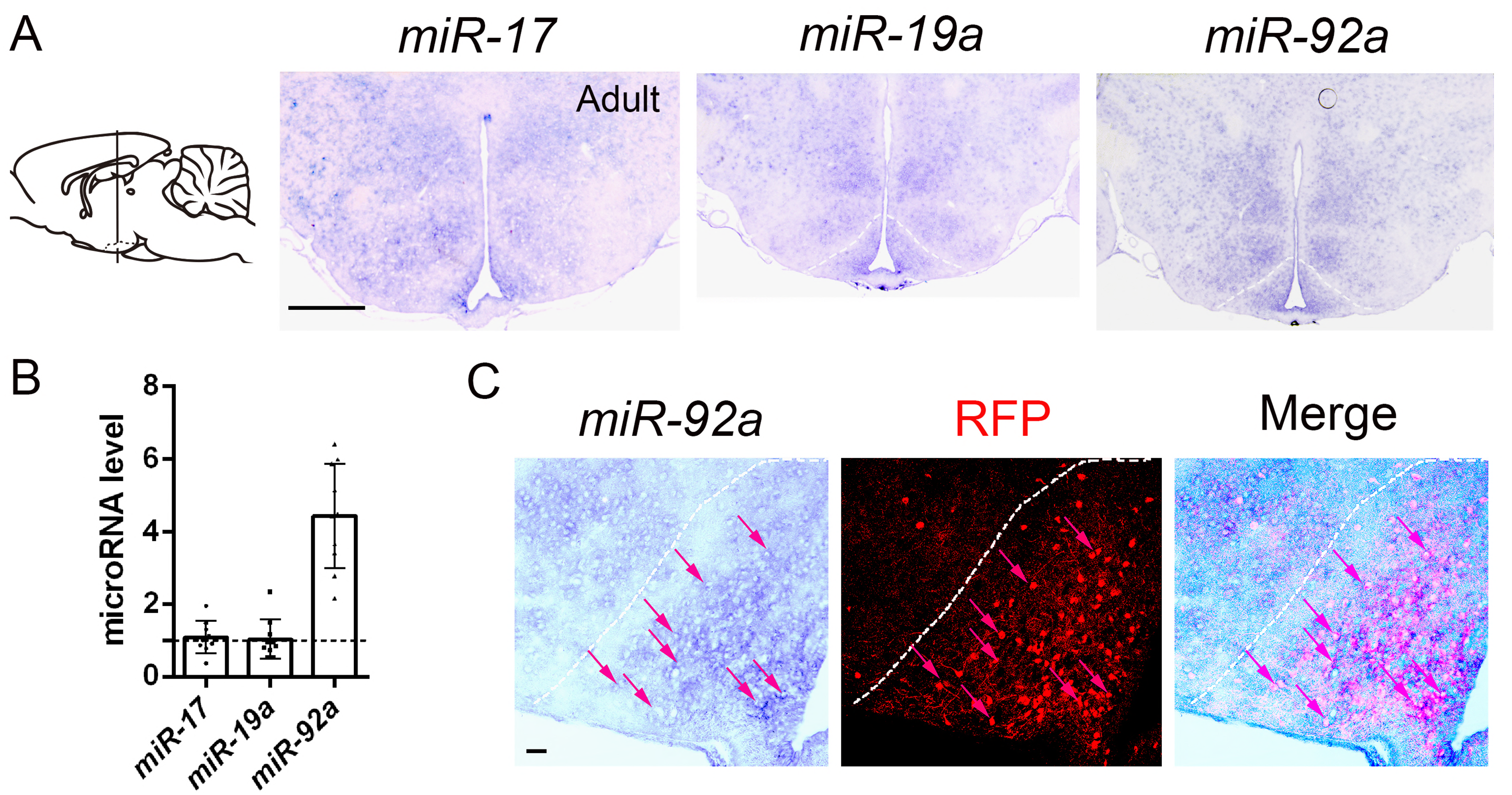
*miR-17-92* expression in the POMC neurons in adult mouse brains. A miR-17, 19a and 92a expression in the adult ARC detected by *in situ* hybridization. Dashed lines illustrate the ARC. Scale Bar: 500μm. B Relative expression level of members in the *miR-17-92* family in the adult ARC detected by real time qRT-PCR. n=9-10 brains in each group. C miR-92a was expressed in most POMC neurons in the ARC. Images of miR-92a *in situ* hybridization (left), RFP immunohistochemistry (middle) and merge of both (right). Dashed lines illustrate the ARC, and arrows mark neurons with co-expression of miR-92a and RFP. Scale Bar: 75μm.

We further examined miR-92a expression in POMC neurons using the *Pomc-Cre;tdTomato* line. In brain sections of *Pomc-Cre;tdTomato* mice, *in situ* hybridization was conducted using the miR-92a LNA probe, followed by immunohistochemistry using anti-RFP antibodies to label POMC neurons. Images of RFP signals and miR-92a expression in the ARC were collected under the fluorescent and bright fields, respectively, and overlaid. Majority of cells (88.2% co-marked cells in RFP positive cells, based on 3 individual brain sections) in the ARC was co-marked with RFP and miR-92a probes (Figure 3C). These results imply that miR-92a might be a major player in miR-17-92 and regulates ARC development.

### Distinct body weight phenotypes in male and female *miR-17-92* KO mice under high fat diet

Considering that miR-17-92 is expressed in POMC neurons in the ARC, we suspected that miR-17-92 might also play a role in regulating body weights. Floxed miR-17-92 transgenic mice (*miR-17-92^flox/flox^*) were bred with *Pomc-Cre* mice to generate conditional *miR-17-92* knockout mice in which miR-17-92 is only deleted in POMC neurons, named *miR-17-92* KO (Bian et al, 2013a; Jin et al, 2016).

We first examined the number of POMC neurons in *miR-17-92* KO mice. *Pomc-Cre;miR-17-92^flox/flox^;tdTomato* mice (called *miR-17-92 KO^tdTomato^*) and *Pomc-Cre*; *miR-17-92^flox/+^;tdTomato* mice (named *Ctrl^tdTomato^*) were generated by breeding *miR-17-92^flox/flox^* mice with *Pomc-Cre;miR-17-92^flox/+^;tdTomato* mice. Immunohistochemistry using anti-RFP antibodies to label POMC neurons was conducted and the total numbers of RFP^+^ neurons were counted in every 5 sections from the anterior to posterior ARC in P0 brains. The number of RFP^+^ POMC neurons did not show significant changes between *miR-17-92* KO and control mice (Figure S2C, D). These results indicate that *miR-17-92* knockout doesn’t affect numbers of POMC neurons.

Similar to *miR-7* knockdown mice, under normal chow diet, *miR-17-92* KO mice did not show significant changes in body weights between control and *miR-17-92* KO mice in either males or females (Figure S5A). Interestingly, high fat diet affected body weights in both male and female *miR-17-92* KO mice, with male *miR-17-92* KO mice heavier than male controls, and female *miR-17-92* KO mice lighter than female controls, when the amount of food intake was similar between control and *miR-17-92* KO mice (Figure 4A and B, and Figure S5B). The body weight change occurred earlier in male than female *miR-17-92* KO mice (displayed weight gain at 15 weeks and weight loss at 20 weeks, respectively) (Figure 4A and B).

**Figure 4.**
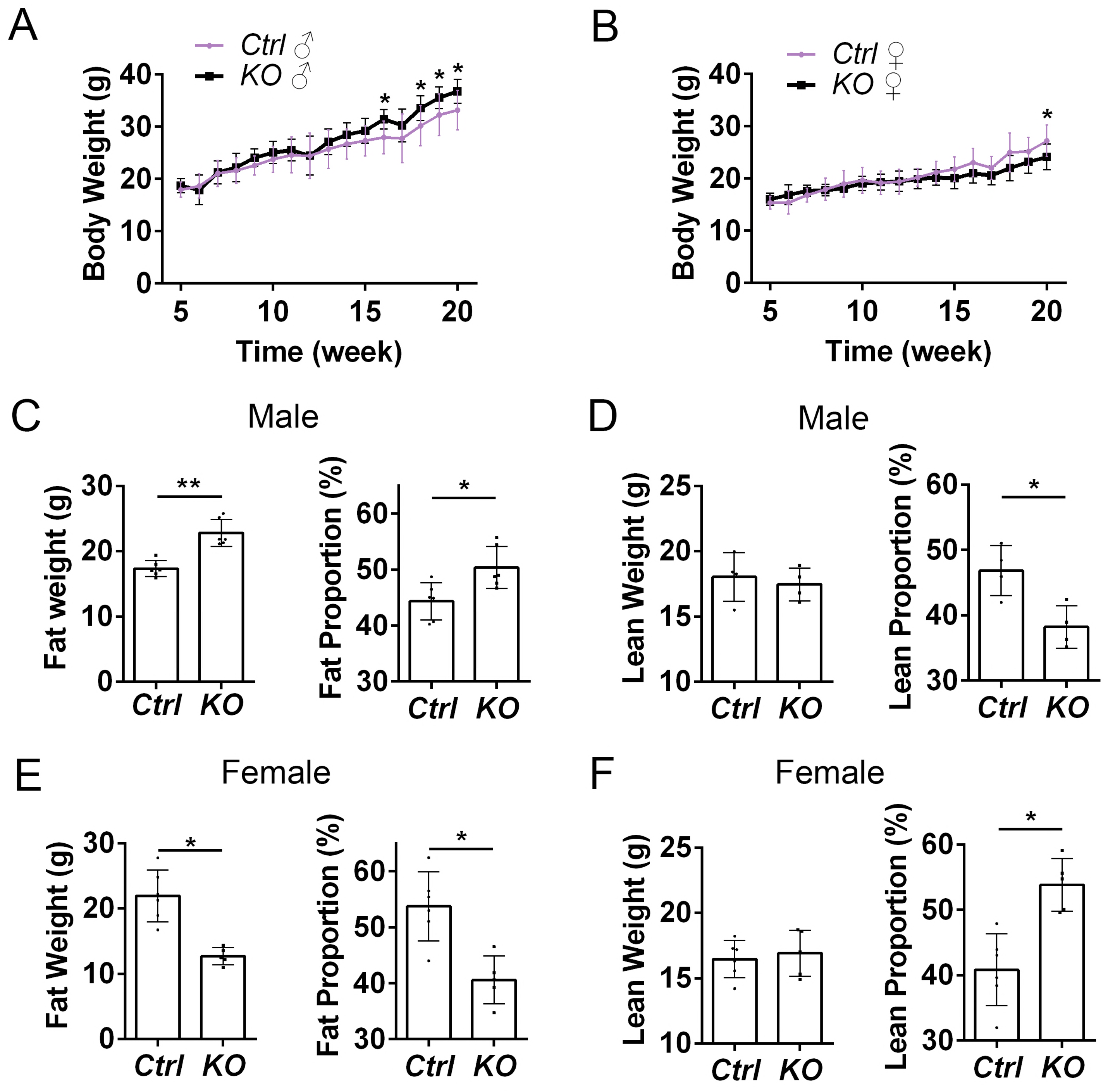
Distinct body weight phenotypes in male and female *miR-17-92* knockout (KO) mice under high fat diet (HFD). A, B Body weight of male (A) and female (B) *miR-17-92* KO and control (Ctrl) mice from 5 to 20 weeks old after treating with HFD. Compared to the same sex control mice, body weights of male *miR-17-92* KO mice were significantly increased. n=6-14 mice at each time point. C, D Body composition of HFD-fed male mice at 25 weeks old. Fat mass and fat proportion (fat weight/body weight) of male *miR-17-92* KO mice were significantly higher than those of controls. Lean mass had no difference between male *miR-17-92* KO and controls, but lean proportion of *miR-17-92* KO mice was significantly lower than that of controls. n=4-6 mice in each group. E, F Body composition of HFD-fed female mice at 25 weeks old. Fat mass and fat proportion (fat weight/body weight) of female *miR-17-92* KO mice were significantly lower than those of controls. Lean mass had no difference between female *miR-17-92* KO and controls, but lean proportion of *miR-17-92* KO mice was significantly higher than that of controls. n=5-6 mice in each group. Values shown are means ± s.e.m. *: P<0.05; **: P<0.01, Student’s t-test.

Moreover, the fat weight and proportion were higher in male *miR-17-92* KO mice than those in male controls, and the lean proportion was less in male *miR-17-92* KO mice than those in male controls (Figure 4C and D). Oppositely, the fat weight and proportion were less in female *miR-17-92* KO mice than those in female controls, and the lean proportion was higher in female *miR-17-92* KO mice than those in female controls (Figure 4E and F). These results indicate that high fat diet affects both males and females when miR-17-92 is deleted in POMC neurons, with gain-of-weight in male KO and loss-of-weight in female KO.

Furthermore, unlike miR-7 knockdown mice, fasting for 48 hours did not affect body weights in either male or female *miR-17-92* KO mice (Figure S5C and D). Additionally, in the IPGTT, blood levels after glucose injection showed no significant changes between *miR-17-92* KO and control mice, no matter males or females, under high fat diet (Figure S5E and F). These data suggest that miR-17-92 knockout does not change capacity of glucose clearance and glucose metabolism.

### Knockdown of miR-7 affects gene expression in the ARC of *miR-7-sp* females

We found that *miR-7-sp* females displayed obvious obesity (Figure 2). To address the molecular basis of miR-7 regulation in this process, total RNA was extracted from the ARC tissues dissected from adult female control and *miR-7-sp* mice (4 months old). RNA sequencing was conducted to screen genes with altered expression in the ARC of female *miR-7-sp* mice. Based on sequencing results, genes with changes of expression levels more than 30% were termed as differentially expressed genes (DEG). Among 1998 DEGs, 1134 genes were upregulated in the ARC of *miR-7-sp* females, while 864 genes were downregulated (Figure S6A).

According to Gene Ontology (GO) analysis (https://david.ncifcrf.gov) of 1134 upregulated genes, genes functioning in “transport” and “regulation of transcription” in the biological process (BP) category, and genes functioning in “nucleic acid binding” and “transcription factor activity” in the molecular function (MF) category, showed higher expression changes than others (Figure S6B). Moreover, in a Kyoto Encyclopedia of Genes and Genomes (KEGG) pathway analysis, genes involved in “alcoholism” and “FoxO signaling pathway” showed higher expression changes (Figure S6C-E).

Mature miRNAs mainly silence target genes through complementary sequence binding in the 3’UTR (Pasquinelli, 2012). Because of the silencing effect of miRNAs, upregulated genes in the ARC of female *miR-7-sp* brains are likely potential targets for miR-7. Among 1134 upregulated genes, 332 genes were predicted as targets of miR-7 by using the algorithm TargetScanMouse 7.1. In the GO analysis of BP, 47 potential targets were identified functioning in mRNA transcription regulation (*P*=9.9E-4) (Figure 5A and C). Real time qRT-PCR further validated differential expression of these genes such as *Foxo1*, *Prkaa1*, *Prkaa2*, *Smad5* and *Ppargc1a* (Figure 5E). Moreover, some of these transcription factors such as Foxo1, Foxo3, Prkaa1, Prkaa2 and Ppargc1a were predicted to form a functional network using a tool in the website String (https://string-db.org/). These results suggest that miR-7 regulates a network of target genes in the POMC neurons.

**Figure 5.**
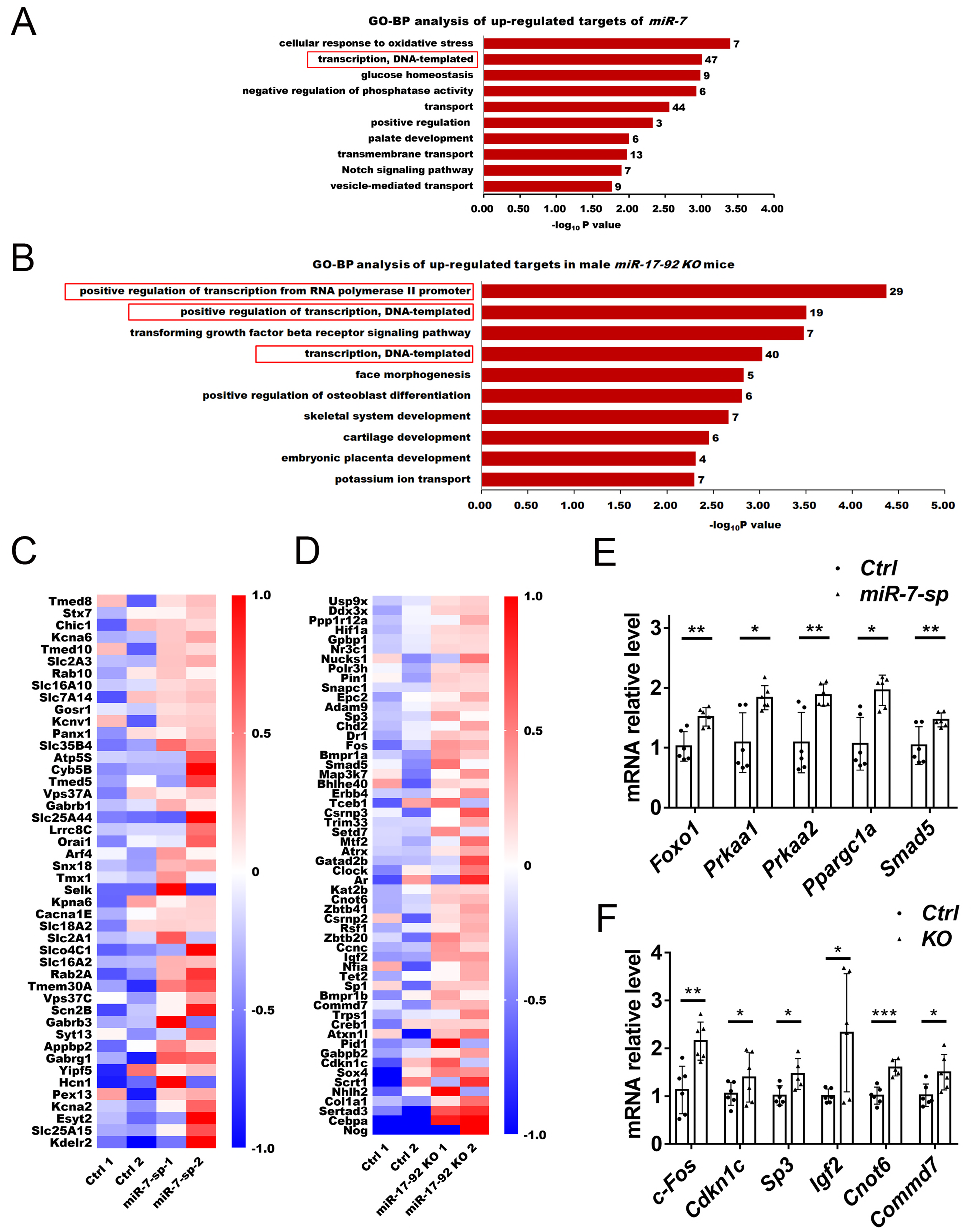
Gene expression changes in the ARC in miR-7-sp and *miR-17-92* KO mice. A The biological process category of gene ontology enrichment analysis (GO-BP analysis) for upregulated target genes for miR-7 in the ARC of female *miR-7-sp* mice. B GO-BP analysis for upregulated target genes for miR-92a in the ARC of male *miR-17-92* KO mice. C Gene expression levels of the “transcription, DNA-templated” group in the GO-BP analysis of female *miR-7-sp* mice. Gene expression levels were calculated by changed fragments per kilobase of transcript per million mapped reads (FPKM) in RNA sequencing data using the formula: [FPKM (sample)-FPKM (average)]/FPKM (average). D Gene expression levels in groups of “positive regulation of transcription from RNA polymerase II promoter”, “positive regulation of transcription, DNA-templated” and “transcription, DNA-templated” in the GO-BP analysis of male *miR-17-92* KO mice. E Relative expression level of *Foxo1, Prkaa1, Prkaa2, Ppargc1a* and *Smad5* in the ARC of female *miR-7-sp* and control mice detected by real time qRT-PCR. F Relative expression level of *c-Fos, Cdkn1c, Sp3, Igf2, Cnot6* and *Commd7* in the ARC of male miR-17-92 *KO* and control mice detected by real time qRT-PCR. Student’s t-test. n=5-6 for each group. Values shown are means ± s.e.m. *: P<0.05; **: P<0.01 and ***: P<0.001, Student’s t-test.

Furthermore, among 864 downregulated genes, 376 genes were grouped in the phosphoproteins and 166 in the acetylation proteins in the keyword features (https://david.ncifcrf.gov) (Figure S6F). Moreover, genes involved in systemic lupus erythematosus (*P*=1.77E-05) and neuroactive ligand-receptor interaction (*P*=2.08E-04) were detected in the KEGG pathway analysis (Figure S6F). These analyses suggest that downregulated genes in the ARC of *miR-7-sp* females might be indirectly regulated by miR-7 and participate in regulation of development and function of POMC neurons.

### Gene expression changes in the ARC in *miR-17-92* knockout mice

Because male *miR-17-92* KO mice displayed heavier body weights than male controls (Figure 4), RNA sequencing analysis was performed using total RNAs extracted from the ARC of adult male control and *miR-17-92* KO mice. Among 3053 DEGs, 2003 genes were upregulated, and 1050 genes were downregulated in the ARC of *miR-17-92* KO males (Figure S7A).

Among upregulated genes, 270 genes were grouped into “regulation of transcription” in the GO-BP category, and RNA binding genes were detected in the GO-MF analysis (Figure S7B). Additionally, in the KEGG pathway analysis, genes involved in “ribosome”, “Huntington’s disease” and “Parkinson’s disease” showed higher expression changes (Figure S7B and C).

Because miR-92a showed higher expression in the ARC (Figure 3), among 2003 upregulated genes, targets for miR-92a were predicted using the TargetScanMouse 7.1 algorithm, and 275 genes were detected. 56 potential target genes that function in transcription-related regulation were detected using the GO-BP analysis (Figure 5B and D). Real time qRT-PCR further validated differential expression of these genes such as *c-Fos, Cdkn1c, Sp3, Igf2, Cnot6* and *Commd7* (Figure 5F).

Moreover, of 1050 downregulated genes, genes involved in “nervous system development” (38 genes) and “homophilic cell adhesion” (25 genes) were detected in the GO-BP analysis, and the gene group of “alcoholism” was the largest group detected in the KEGG pathway analysis (Figure S7D). These data suggest that deletion of miR-17-92, similar to miR-7 knockdown, indirectly affects expression of genes regulating neuronal development.

### Upregulation of female low-expression genes is associated with female-specific obesity in *miR-7-sp* mice

Because male and female *miR-7-sp* and *miR-17-92* KO mice showed differential response in body weight under high fat diet (Figures 2 and 4), we speculated that gene expression in the ARC might display sex-specific differences. To test this idea, RNA sequencing analysis was performed using total RNAs extracted from the ARCs of wildtype adult male and female mouse brains. Among differentially expressed genes in the male and female ARCs (male versus female), genes with fold changes less than 0.77 were considered as male low-expression genes (i.e. female high-expression genes), while genes with fold changes more than 1.3 were considered as female low-expression genes (i.e. male high-expression genes).

In the male and female ARCs, while 10,478 genes showed no differential expression between males and females, 2957 genes displayed sex-biased expression, with 1641 genes showing male low-expression and 1316 genes showing female low-expression (Figure 6A). Real time qRT-PCR further validated differential expression of these genes such as *Foxo1, Fbxo45* and *Rybp* in the ARC of wildtype brains (Figure 6B). Moreover, the GO-BP analysis of 2957 sex-differentially expressed genes (SDEGs) detected 409 genes (about 13.8%) that are involved in “regulation of transcription”, suggesting a role of gene expression regulation of SDEGs (Figure S8).

**Figure 6.**
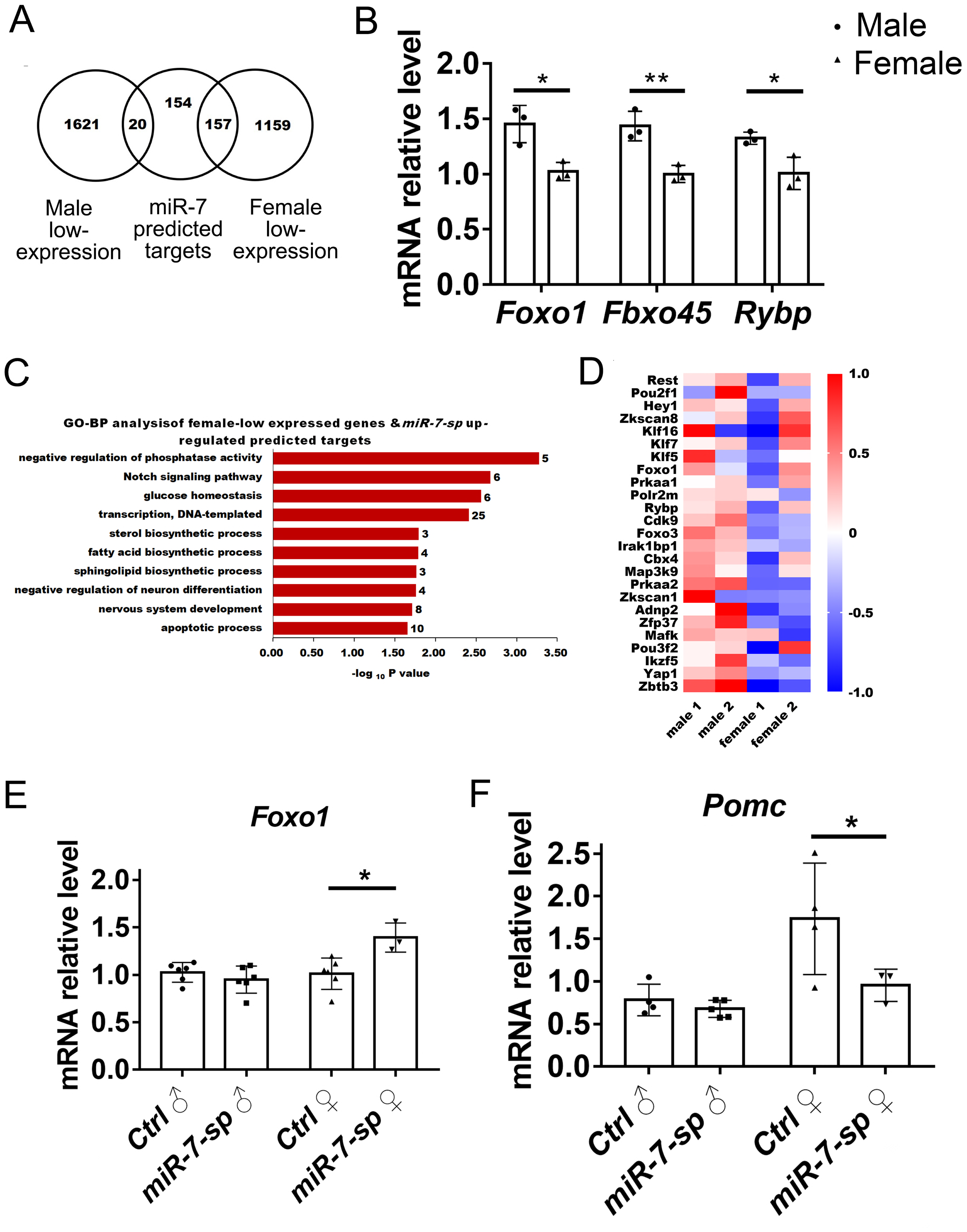
Upregulation of female low-expression genes is associated with female-specific obesity in miR-7-sp mice. A Venn diagram of sex-differentially expressed genes and upregulated predicted targets for miR-7. B Validation of female low-expressed genes *Foxo1*, *Fbxo45* and *Rybp* in the ARC of wildtype mouse brains detected by real time qRT-PCR. n=3 brains for each group. C Among 331 predicted targets for miR-7, 20 genes showed male low-expression, and 157 genes displayed female low-expression. A diagram of the biological process category of gene ontology enrichment analysis (GO-BP analysis) of female low-expressed genes. D Gene expression levels of the “transcription, DNA-templated” group in the GO-BP analysis of the 157 genes. E The relative expression level of *Foxo1* in the ARC of HFD-fed *miR-7-sp* and control mice detected by real time qRT-PCR. n=3-6 brains for each group. F The relative expression level of *Pomc* in the ARC of HFD-fed *miR-7-sp* and control mice detected by real time qRT-PCR. n=3-5 brains for each group. Values shown are means ± s.e.m. *: P<0.05; **: P<0.01, Student’s t-test (D) and ANOVA test (E and F).

Given that female *miR-7-sp* mice displayed obvious obesity, we explored whether upregulated and predicted target genes for miR-7 in the ARC of *miR-7-sp* brains show sex-differential expression. Interestingly, among 331 predicted targets, 20 genes showed male low-expression, i.e. female high-expression, while 157 potential target genes displayed female low-expression, i.e. male high-expression (Figure 6A). These data suggest that miR-7 knockdown causes upregulation of genes that normally have low expression in the female ARC. Moreover, the GO-BP analysis based on 157 female low-expression genes identified top 10 categories, and 25 genes functioning in regulating transcription (Figure 6C and D).

Furthermore, to examine whether female low-expression genes respond to HFD treatment, we quantified *Foxo1* expression level by qRT-PCR in male and female *miR-7-sp* and control mice fed with HFD. We found elevated *Foxo1* expression in the ARC of female *miR-7-sp* mice, compared to their same sex controls, while there were no detectable changes in the ARC of male *miR-7-sp* and their same sex controls (Figure 6E). In addition, it has been shown that Foxo1 can repress Pomc level in POMC neurons (Ma et al, 2015), so we quantified *Pomc* expression level by qRT-PCR. While there was no difference in male *miR-7-sp* and control mice, there was a significant reduction of *Pomc* expression in the ARC of female *miR-7-sp* mice, compared to their same sex controls, upon HFD treatment (Figure 6F).

Taken together, upregulation of genes that normally show low expression in ARCs of females might contribute to female-specific obesity when miR-7 is knocked down.

### *miR-17-92* KO specifically affects sex-differentially expressed genes in the ARC

Because female and male *miR-17-92* KO mice showed distinct responses towards HFD (Figure 4), we explored whether predicted target genes for miR-92a among upregulated genes in the ARC of *miR-17-92* KO brains show sex-differential expression. Among 274 predicted targets, 111 genes showed male low-expression, and 14 genes displayed male high-expression (Figure 7A). These results suggest that *miR-17-92* knockout results in upregulation of genes that normally have low expression in the male ARC. Real time qRT-PCR further validated differential expression of these genes such as *Ddx3x, Bmpr1A* and *Clock* in the ARC of wildtype mouse brains (Figure 7B). Moreover, GO-BP analysis of 111 male low-expression genes identified 22 genes that are involved in transcription regulation (Figure 7C and D).

**Figure 7.**
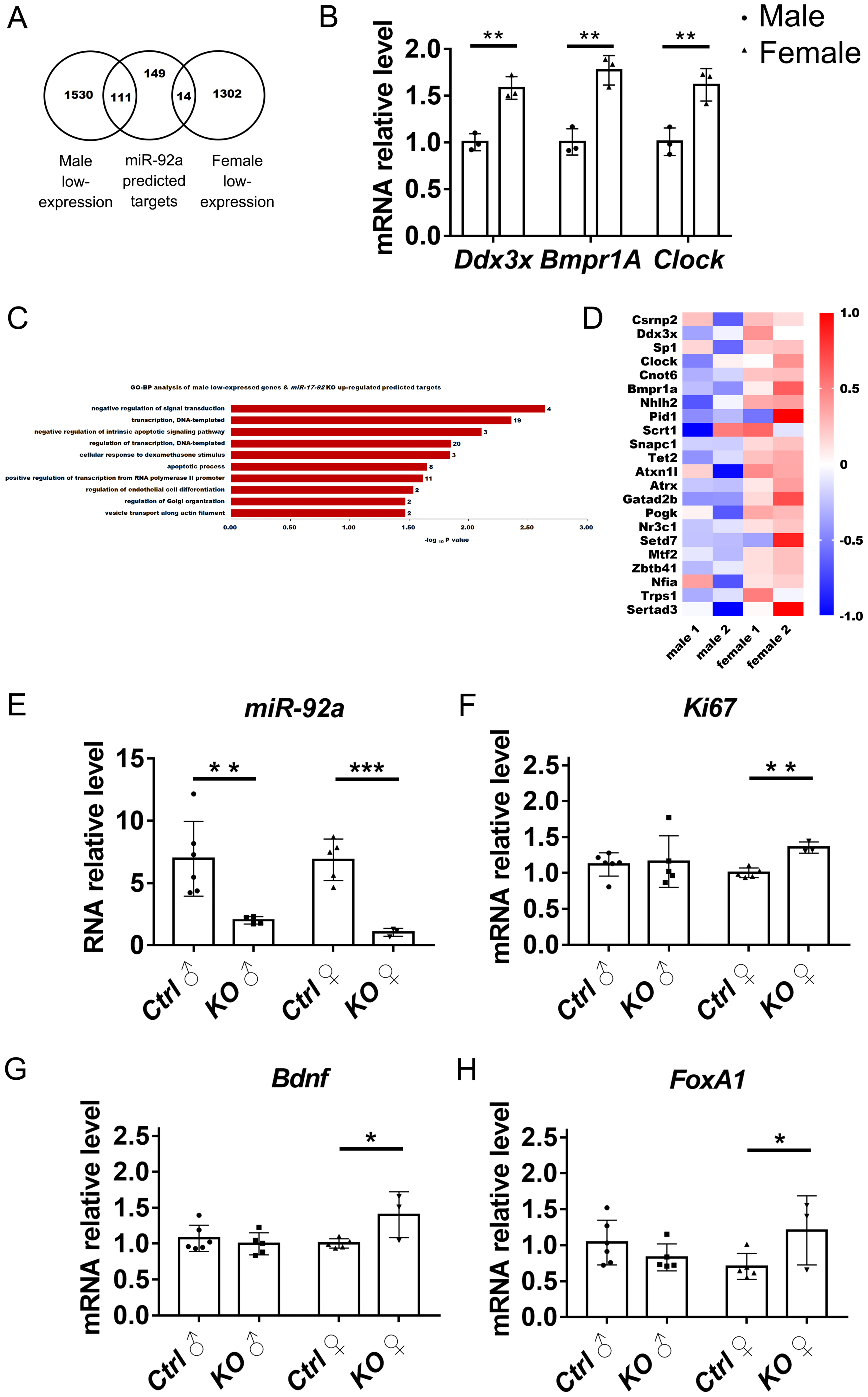
*miR-17-92* knockout (KO) specifically affects sex-differentially expressed genes in the ARC. A Venn diagram of sex-differentially expressed genes and upregulated predicted targets for miR-92. B Validation of male low-expressed genes *Ddx3x, Bmpr1A* and *Clock* in the ARC of wildtype mouse brains detected by real time qRT-PCR. n=3 brains for each group. C Among 274 predicted targets for miR-92, 111 genes showed male low-expression, and 14 genes displayed male high-expression. A diagram of the biological process category of gene ontology enrichment analysis (GO-BP analysis) of male low-expressed genes. D Gene expression levels of the “transcription, DNA-templated”, “regulation of transcription, DNA-templated” and “positive regulation of transcription from RNA polymerase II promoter” groups in the GO-BP analysis of the 111 genes. E-H The relative expression level of *miR-92a, Bdnf, ki67* and *FoxA1* in the ARC of HFD-fed *miR-17-92* KO and control mice detected by real time qRT-PCR. n=3-6 brains for each group. Values shown are means ± s.e.m. *: P<0.05; **: P<0.01 and ***: P<0.001, Student’s t-test (D) and ANOVA test (E-H).

Furthermore, we identified significant reduction of miR-92a expression in the ARC of *miR-17-92* KO mice compared to control mice, which were both fed with HFD, using qRT-PCR (Figure 7E). We then examined expression levels of sex-differentially expressed genes that are associated with neurogenesis such as *Ki67* and *Bdnf* in the ARC of *miR-17-92* KO and control mice upon HFD treatment. While there was no difference in male *miR-17-92* KO and control mice, there was a significant increase of *Ki67* and *Bdnf* expression in the ARC of female *miR-17-92* KO mice, compared to their same sex controls (Figure 7F and G). Moreover, FoxA1 has been shown to act as a key factor for both androgen receptor (AR) and estrogen receptors (ER) to mediate sexual dimorphism (Hurtado et al, 2011; Li et al, 2012; Reddy et al, 2015; Sahu et al, 2011; Yang & Yu, 2015), we thus further measured its expression. We found elevated *FoxA1* expression in the ARC of female *miR-17-92* KO mice, compared to their same sex controls, while there were no detectable changes in the ARC of male *miR-17-92* KO and their same sex controls (Figure 7H).

In summary, these data suggest that sex-specific responses to HFD in *miR-17-92* KO might be due to distinct altered expression of genes which normally show sex differential expression in the ARC.

## Discussion

Energy expenditure and obesity are regulated by hypothalamus, in particular, the ARC that consists of POMC neurons. Studies have shown that POMC neurons modulate sex-specific changes of body weights in diet-induced obesity (Burke et al, 2016; Medrikova et al, 2012; Salinero et al, 2018). Here we show that POMC neuron-specific miR-7 knockdown and *miR-17-92* knockout mice display sex-different response to high fat diet. We further demonstrate that altered expression of genes that normally show high or low expression levels in the ARC of male or female mice are associated with sex-specific obesity.

Accumulating evidence has shown important roles of miRNAs in regulating body weights. For example, long-time systemic inhibition of miR-21 can relieve obesity in leptin receptor-deficient (*db/db*) mice by decreasing the adipocyte size and serum triglycerides (Seeger et al, 2014), and *miR-155* knockout specifically protects high fat diet induced obesity in female mice by increasing energy expenditure (Gaudet et al, 2016). Furthermore, miR-219 conditional knockdown in the ventromedial hypothalamus (VMH), which also participates in obesity and energy expenditure, increases the risk of obesity, while miR-219 overexpression relieves diet-induced obesity (Schroeder et al, 2018). Our study here shows that miR-7 and miR-17-92 regulate body weight, glucose consumption and diet-induced obesity in a sex preference manner.

MiRNA miR-7 is crucial for neural development and function. Knocking down miR-7 using the miRNA sponge specifically in the cortex causes a smaller brain due to altered neural progenitor proliferation and differentiation (Pollock et al, 2014). Studies have shown that miR-7 binds to circular RNA *Cdr1as* to modulate brain functions (Piwecka et al, 2017). Moreover, the miR-17-92 cluster plays essential roles in controlling expansion of neural stem cells and neural progenitors, and in turn the brain size (Bian et al, 2013a; Garg et al, 2013). miR-17-92 also regulates mood associated behaviors by maintaining adult neural stem cell proliferation (Jin et al, 2016). Here we show that both miR-7 and miR-17-92 play a role in regulating ARC-related body weight control. We have found that knocking down miR-7 and knocking out miR-17-92 specifically in the ARC don’t affect the number of POMC neurons, suggesting distinct role of these miRNAs in the cortex and hypothalamus. These results also imply that altered gene expression by miRNAs in POMC neurons, instead of the number of them, is crucial for hypothalamus function. Furthermore, miR-7 knockdown and miR-17-92 knockout mice don’t show body weight changes under normal chow diet, while they display obesity under high fat diet. Our results support the perspective view of miRNA functions, while miRNAs play robust roles in developmental processes, they also excel functions in a tissue-specific manner, in particular under stress conditions (Ebert & Sharp, 2012; Leung & Sharp, 2010).

Interestingly, we have found that only female miR-7 knockdown mice and male *miR-17-92* knockout mice display diet-induced obesity. Previous reports have shown sex-specific changes of body weights, which is regulated by POMC neurons (Burke et al, 2016; Medrikova et al, 2012; Salinero et al, 2018). However, how POMC neurons modulate sex-specific changes of body weight is unclear. Here we have found that some genes display differential expression levels in the male and female ARCs in wildtype mouse brains. How sex-differential expression of these genes are established, and whether their expressions are maintained by hormone or other factors are still unclear (Burke et al, 2016; Plum et al, 2007). Because miRNAs normally function through silencing target genes, genes upregulated in the ARCs of miR-7 knockdown and miR-17-92 knockout brains are likely potential targets for these miRNAs. Interestingly, we have found that some of these upregulated target genes are enriched in sex-differentially expressed genes. In particular, miR-7 target genes, which normally show low-expression in wildtype female ARCs, are upregulated in female ARCs of miR-7-sp mice, while miR-92 target genes, which normally display low-expression in wildtype male ARCs, are upregulated in male ARCs of *miR-17-92* knockout mice. The gene expression profile shift of sex-differentially expressed genes in male or female ARCs, caused by altered miR-7 and miR-17-92 expression, might contribute to sex-specific body weight changes. The future study should examine why and how female and male ARCs display gene expression changes upon miR-7 knockdown and miR-17-92 knockout, respectively.

Because of the feature of miRNAs in silencing target genes, it is likely that sex-specific body weight changes in miR-7 knockdown and *miR-17-92* knockout mice are caused by altered targets for miR-7 and miR-17-92. Studies have shown that decreased Pomc level can induce obesity (Challis et al, 2004; Greenman et al, 2013). And the Pomc level is usually repressed by Foxo1 (Ma et al, 2015). We have found elevated expression of *Foxo1* in the female not male ARC in miR-7 knockdown brains, which implies potential regulatory consequence Foxo1, Pomc level and body weight changes. Furthermore, FoxA1 has been reported to mediate sexual dimorphism (Li et al, 2012; Reddy et al, 2015). We here have found increased *FoxA1* level in the female not male ARC in *miR-17-92* knockout brains. Moreover, because one miRNA can have many targets, it is likely that sex-specific body weight changes are a result of alteration of multiple gene pathways that are normally repressed by miR-7 and miR-17-92. Our RNA-seq data of a group of sex-differentially expressed genes also support this possibility. The future work should dissect major miRNA targets that mediate sex-specific functions of hypothalamus.

Taking together, our studies have shown that miR-7 and miR-17-92 play crucial roles in regulating body weight and glucose consumption, which are mediated by POMC neurons in the ARC of hypothalamus. Importantly, miR-7- and miR-17-92-mediated body weight control upon high fat diet treatment displays sex-specific feature, which is likely caused by altered expression of sex-differential genes in the ARC. Our study should shed new light on mechanistic and preventive investigation of obesity in the future.

## Acknowledgements

We thank members of the Sun laboratory for their valuable discussions and advice. This work was supported by an R01-MH083680 grant from the NIH/NIMH (T. S.), and the National Natural Science Foundation of China (81471152, 31771141 and 81701132).

## Author contributions

Conceived and designed the experiments: **Y.G**. and **T.S**. Experiment: **Y.G**., **Z.Z**. and **A. P**. Result analysis: **Y.G**., **J.L**. and **R.Z**. Wrote the paper: **Y.G**. and **T.S**. Edited the paper: **T.S**.

